# Male-driven reinforcement and cascade reinforcement in darters

**DOI:** 10.1101/171231

**Authors:** R. L. Moran, R. C. Fuller

## Abstract

Reinforcement can act in response to maladaptive hybridization by selecting for increased discrimination against heterospecifics mates in sympatry compared to allopatry (i.e., reproductive character displacement - RCD). Additionally, reinforcement can select for increased discrimination against heterospecifics in a fighting context in sympatry compared to allopatry (i.e., agonistic character displacement - ACD). Because it directly affects conspecific recognition traits (signals and/or preferences), reinforcement between two species in sympatry can incidentally cause behavioral isolation among populations within a species, termed cascade reinforcement. Here we demonstrate that a pattern consistent with male-driven RCD and ACD is present between two groups of darters (orangethroat darter clade *Ceasia* and rainbow darter *Etheostoma caeruleum*). Increased male discrimination against heterospecific females as mates and increased male discrimination against heterospecific males as competitors is present in sympatry. Furthermore, there is a pattern consistent with male-driven cascade RCD and cascade ACD within *Ceasia*. We found low levels of discrimination between two species of *Ceasia* that occur in allopatry from one another and in allopatry with *E. caeruleum*. This result contrasts that of a recent study which observed high levels of behavioral isolation between *Ceasia* species that occurred in sympatry with *E. caeruleum*. We suggest reinforcement between *Ceasia* and *E. caeruleum* in sympatry has led to the evolution of behavioral isolation between lineages within *Ceasia* that occur in sympatry with *E. caeruleum* but in allopatry with respect to one another (i.e., cascade reinforcement). This study demonstrates the ability of male behavior to simultaneously drive sympatric and allopatric speciation via reinforcement.

## Introduction

Reinforcement is unique among speciation mechanisms in that it can directly select for increased behavioral isolation between two species in response to the production of unfit hybrids in areas of sympatry (Dobzhansky 1940; Butlin 1987; Kelly and Noor 1996). Consequently, reinforcement causes mating traits (signals and/or preferences) to diverge between species in sympatry (but not allopatry), leading to a pattern of reproductive character displacement (RCD; Brown and Wilson 1956; Howard 1993; Coyne and Orr 2004). RCD is the classic signature used to detect reinforcement, and is evidenced by increased discrimination against heterospecific mates in sympatry compared to allopatry (Servedio and Noor 2003).

Although the most recognized result of reinforcement is increased behavioral isolation between two species in sympatry, recent research has suggested that reinforcement can also initiate allopatric speciation (Ortiz-Barrientos et al. 2009; Pfennig and Pfennig 2009; Hoskin and Higgie 2010). By directly affecting traits associated with behavioral isolation with a closely related sympatric species, reinforcement may alter behavioral isolation among populations within a species. Heightened behavioral isolation among populations of species that also experience reinforcement with a close relative has been documented in numerous empirical examples across a variety of taxa (e.g., Nosil et al. 2003; Hoskin et al. 2005; Higgie and Blows 2007, 2008; Lemmon 2009; Porretta and Urbanelli 2012; Bewick and Dyer 2014; Pfennig and Rice 2014; Kozak et al. 2015). When reinforcement occurs independently in isolated populations throughout a species range, stochastic processes may cause different mating traits underlying behavioral isolation to diverge in different populations (reviewed in Comeault and Matute 2016). This can cause behavioral isolation to evolve among allopatric populations within a species, termed cascade reinforcement (Ortiz-Barrientos et al. 2009).

To illustrate this concept, we show a hypothetical range map with one wide-ranging species (A) and several populations/newly formed species (B1-B4) (Fig. 1). Populations B1-B4 are all allopatric to one another. Populations B1 and B2 are sympatric with respect to A, but populations B3 and B4 are allopatric with respect to species A. In this scenario, cascade reinforcement predicts that there is reinforcement between A and B1 and between A and B2, independently. Behavioral isolation is thus heightened between A and B1 and between A and B2. However, because slightly different traits have evolved in B1 and B2 in response to reinforcing selection, behavioral isolation is also heightened between B1 and B2 as a *by-product* of reinforcement. In contrast, there is no reinforcement between A and B3 or A and B4, and behavioral isolation is low in these pairs of taxa. Likewise, behavioral isolation is low between B3 and B4. The critical test for cascade reinforcement is whether allopatric populations that experience reinforcement with a more distant relative (i.e., B1 and B2) have higher behavioral isolation than allopatric populations that do not experience reinforcement (i.e., B3 and B4).

**Figure 1.**
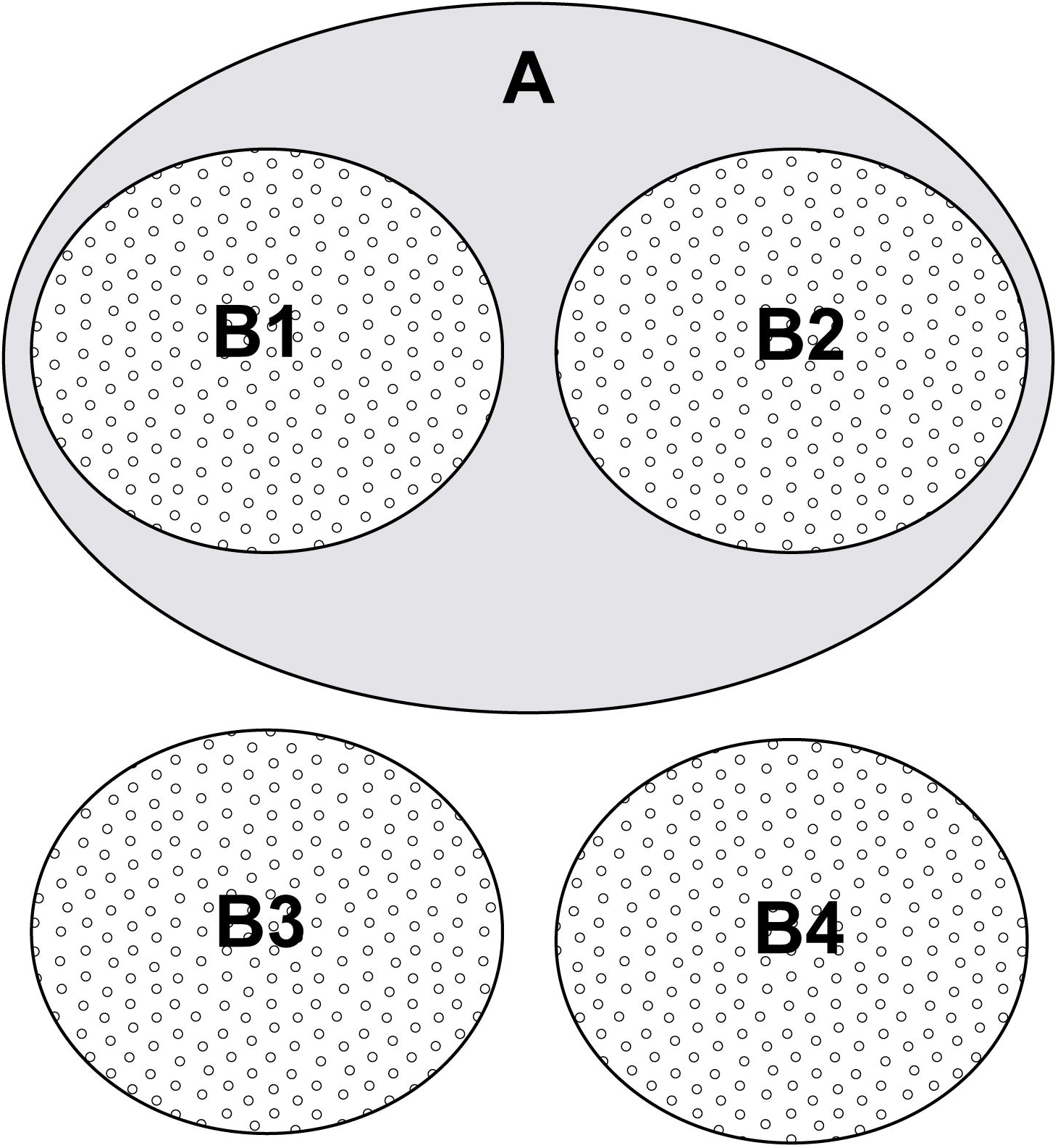
Hypothetical ranges for two species, A and B, with four populations shown within B (B1-B4). Populations B1-B4 are allopatric from one another. Populations B1 and B2 are sympatric with species A. Populations B3 and B4 are allopatric from species A. Reinforcement occurring independently between A and B1 and between A and B2 can incidentally cause heightened behavioral isolation between B1 and B2 compared to behavioral isolation present between B3 and B4 (i.e., cascade reinforcement).

The term reinforcement has primarily been used to describe selection acting against maladaptive heterospecific mating, but reinforcement can also act via selection against maladaptive heterospecific fighting over resources (such as mates). This form of reinforcement can cause aggressive signals and/or recognition of competitor’s signals to diverge between species in sympatry, resulting in a pattern of agonistic character displacement (ACD; Grether et al. 2009; Okamoto and Grether 2013). A pattern of ACD is said to be present when males from two different species are less likely to engage in contests when they occur in sympatry compared to allopatry with one another. Both reinforcement leading to RCD and reinforcement leading to ACD may contribute to speciation. Although numerous studies have shown that RCD can occur among populations within species due to cascade reinforcement (e.g., Nosil et al. 2003; Hoskin et al. 2005; Lemmon 2009; Rice and Pfennig 2010; Pfennig and Rice 2014; i.e., cascade RCD, hereafter CRCD), whether reinforcement can also secondarily lead to ACD among populations within species (i.e., cascade ACD, hereafter CACD) has yet to be determined.

Distinguishing between RCD and ACD is essential to determining the underlying selective pressure (i.e., selection against heterospecific mating or heterospecific fighting) and relative contribution of male-female and male-male interactions in driving speciation. However, disentangling the relative contributions of RCD and ACD to speciation can be difficult because many sexually selected traits are important in both intersexual contexts (such as female mate choice) and intrasexual contexts (such as male-male competition over mates) (Alatalo et al. 1994; Berglund 1996; Sætre et al. 1997; Dijkstra et al. 2007; Saether et al. 2007; Lackey and Boughman 2013; Tinghitella et al. 2015). Here we examine both RCD and ACD using a system where males discriminate against both heterospecific female mates and heterospecific male rivals, but female mate choice appears to be absent.

This study focusses on two groups of darters in the in the subgenus *Oligocephalus*: the orangethroat darter clade *Ceasia* and the rainbow darter *Etheostoma caeruleum*. The clade *Ceasia* consists of 15 recently diverged species, which all occur in allopatry from one another. Twelve of the 15 *Ceasia* species occur in sympatry with *E. caeruleum* throughout their range (Ceas and Page 1997; Page and Burr 2011). One wide-ranging *Ceasia* species (orangethroat darter *Etheostoma specatbile*), occurs both in sympatry and in allopatry with *E. caeruleum*. The reaming two *Ceasia* species (plains darter *E. pulchellum* and plateau darter *E. squamosum*) are completely allopatric with respect to *E. caeruleum*. *Ceasia* and *E. caeruleum* have similar male nuptial coloration, ecology, and mating behavior (Page 1983; Page and Burr 2011). There is little evidence that male coloration in *Ceasia* and *E. caeruleum* is under selection by females in intersexual mate choice, as females appear to lack preferences within or among species (Pyron 1995; Fuller 2003; Zhou et al. 2015; Moran et al. in press). Instead, male coloration is under intrasexual selection, functioning as an aggressive signal in male-male competition over access to females (Zhou and Fuller 2016; Moran et al. in press).

Evidence from several recent studies suggests that reinforcement is likely occurring in this system. First, hybridization occurs at low rates between *Ceasia* and *E. caeruleum* in nature (Zhou and Fuller 2014; Moran et al. in press). Second, postzygotic isolation is present between at least one *Ceasia* species (*E. spectabile*) and *E. caeruleum*. F1 hybrid clutches between these species have a male-skewed sex ratios (Zhou 2014), and backcross hybrids suffer from dramatically reduced viability (R. Moran unpubl. data). Third, a recent study found high levels of male-driven behavioral isolation between four species of *Ceasia* (strawberry darter *E. fragi*, current darter *E. uniporum*, brook darter *E. burri*, and *E. spectabile*) and their respective sympatric populations of *E. caeruleum* (Moran et al. in press). Fourth, a pattern consistent with RCD was observed in a no-choice mating experiment between *E. spectabile* and *E. caeruleum* (Zhou and Fuller 2014). Zhou and Fuller (2014) found that pairings of allopatric female *E. spectabile* and allopatric male *E. caeruleum* yielded more eggs than pairings of sympatric female *E. spectabile* and sympatric male *E. caeruleum*. Together, the results of these previous studies strongly suggest that reinforcement may be occurring between *Ceasia* and *E. caeruleum*. However, the no-choice assay performed previously by Zhou and Fuller (2014) was not able to measure the contribution of each sex to behavioral isolation in sympatry, and did not consider the role of male competition (an important component of behavioral isolation in darters; Zhou et al. 2015; Martin and Mendelson 2016; Moran et al. in press). Here we provide a direct measure of mating behavior in both sexes to test for reinforcement. We examine whether behavioral isolation is lower (or absent) between *Ceasia* and *E. caeruleum* when they occur in allopatry from one another.

There is also reason to suspect that cascade reinforcement may be present within *Ceasia*. Moran et al. (in press) found that surprisingly high levels of behavioral isolation are present among recently diverged allopatric species of *Ceasia*. Male *Ceasia* discriminate against heterospecific *Ceasia* female mates and against heterospecific *Ceasia* male rivals (Moran et al. in press). The behavioral isolation among *Ceasia* species is not associated with differences in male color pattern or genetic distance. Notably, the *Ceasia* species examined by Moran et al. (in press) all occur sympatrically with *E. caeruleum*. Therefore, the high levels of behavioral isolation observed among *Ceasia* may be due to reinforcement between sympatric *Ceasia* and *E. caeruleum* incidentally causing cascade reinforcement within *Ceasia*. Here we test this hypothesis by examining whether behavioral isolation is higher among *Ceasia* species that are sympatric versus those that are allopatric with *E. caeruleum*.

Our first aim was to determine whether a pattern consistent with RCD and/or ACD is present between *Ceasia* and *E. caeruleum*. To do this, we compared discrimination against heterospecifics in the context of male mate choice, female mate choice, and male aggression among *Ceasia* and *E. caeruleum* populations/species that were sympatric versus allopatric with respect to one another. RCD predicts higher levels of male and/or female discrimination against heterospecific mates in sympatry compared to allopatry. If RCD is present due to alterations in male preferences in sympatry, then males from sympatric populations should only pursue conspecific females, whereas males from allopatric populations should pursue both conspecifics and heterospecifics. Likewise, if RCD is present due to changes in female preferences in sympatry, then females from sympatric populations should only perform nosedigs (a behavior that directly proceeds spawning) when they are being guarded by conspecific males. Allopatric females should not show such discrimination. ACD predicts higher levels of male discrimination against heterospecific male rivals in sympatry compared to allopatry. If ACD is present due to alterations in male competitor recognition in sympatry, then males from sympatric populations should engage in more aggressive interactions with conspecific males compared to heterospecific males, whereas males from allopatric populations should not bias their aggression towards conspecifics.

Our second aim was to determine whether a pattern consistent with CRCD and/or CACD is present within *Ceasia*. To do this, we measured behavioral isolation between *Ceasia* species that occur allopatrically from one another and allopatrically from *E. caeruleum*. We then compared these estimates to previous measures of behavioral isolation among *Ceasia* species that occur sympatrically with *E. caeruleum* (Moran et al in press). CRCD predicts higher levels of mate discrimination against heterospecific *Ceasia* in populations/species that occur in sympatry with *E. caeruleum*. CACD predicts higher levels of male discrimination against heterospecific *Ceasia* male rivals in populations/species that occur in sympatry with *E. caeruleum*.

## Methods

### Mating system details

During the spring spawning season, both *Ceasia* and *E. caeruleum* travel to shallow gravel riffles in headwater streams (Hubbs and Strawn 1957; Hubbs 1985; Heins et al. 1996). Females look for a suitable place to lay eggs by preforming “nosedigs” in which they jab their snout into the gravel. One to several males will swim in tandem along a female as she searches for a spawning location. Males fight aggressively to ward off rival males, either by actively chasing them off or by flaring their dorsal and anal fins in a threat display. When the female is ready to spawn, she will dive into the substrate and position herself so that only her head and caudal fin are fully visible. Fighting amongst males escalates at this point, as they attempt to secure access to the female. During spawning, a male positions himself above the female and they simultaneously release sperm and eggs into the substrate. Spawning often involves multiple males mating simultaneously with one female, and males sometimes exhibit sneaking behavior. Females will ovulate clutches of up to 200 eggs periodically throughout the spawning season, but only release a few eggs per spawning bout (Heins et al. 1996; Fuller 1998). Hence, the female must spawn multiple times to fertilize all the eggs from a given clutch.

### Study species/populations and collection locations

We conducted behavioral trials focusing on two wide-spread species of *Ceasia* (*E. spectabile* and *E. pulchellum*), and *E. caeruleum. E. spectabile* occurs in sympatry with *E. caeruleum* in its eastern range, but it occurs in allopatry with respect to *E. caeruleum* in the western part of its range. *E. pulchellum* is allopatric to *E. caeruleum* throughout its range. All species of *Ceasia* occur in allopatry from one another. Hereafter, when we describe populations of *Ceasia* or *E. caeruleum* as being sympatric or allopatric, we are referring to the geographic relationship between *Ceasia* and *E. caeruleum* (not between *Ceasia* species). We used three *Ceasia* study populations: sympatric *E. spectabile*, allopatric *E. spectabile*, and allopatric *E. pulchellum*. We also used a sympatric *E. caeruleum* population (for comparisons with the sympatric *E. spectabile* population) and an allopatric *E. caeruleum* population (for comparisons with the allopatric *E. spectabile* and the allopatric *E. pulchellum* populations). Table S1 shows collection locations for each of the five darter populations used in this study.

In 2016, we conducted dichotomous male choice assays and male competition assays (detailed below) to measure behavioral isolation in sympatric and allopatric pairings of *Ceasia* and *E. caeruleum* (Table 1). In 2017, we conducted dichotomous male choice assays and male competition assays to measure behavioral isolation between the two allopatric *Ceasia* species (Table 2). We then examined the results of these experiments in combination with data from a previous study (Moran et al. in press) to look for patterns of RCD and ACD between *Ceasia* and *E. caeruleum*, and patterns of CRCD and CACD within *Ceasia*.

**Table 1.** Comparisons made in dichotomous choice trials, which included a focal male *Ceasia* and a conspecific female *Ceasia* with (A) an *E. caeruleum* female, or (B) a heterospecific *Ceasia* female.

| **Focal *Ceasia* male** | **Conspecific female** | **Heterospecific female** | **n** |
| --- | --- | --- | --- |
| Allopatric *E. spectabile* | Allopatric *E. spectabile* | Allopatric *E. caeruleum* | 12 |
| Allopatric *E. pulchellum* | Allopatric *E. pulchellum* | Allopatric *E. caeruleum* | 12 |
| Sympatric *E. spectabile* | Sympatric *E. spectabile* | Sympatric *E. caeruleum* | 12 |

**B. Conspecific *Ceasia* vs. heterospecific *Ceasia* dichotomous male choice trials**
| **Focal *Ceasia* male** | **Conspecific female** | **Heterospecific female** | **n** |
| --- | --- | --- | --- |
| Allopatric *E. spectabile* | Allopatric *E. spectabile* | Allopatric *E. pulchellum* | 12 |
| Allopatric *E. pulchellum* | Allopatric *E. pulchellum* | Allopatric *E. spectabile* | 12 |

**Table 2.** Comparisons made in male competition trials, which included a conspecific focal male and focal female *Ceasia* pair together with (A) a conspecific *Ceasia* rival male, (B) an *E. caeruleum* rival male, or (C) a heterospecific *Ceasia* rival male.

| ***Ceasia* focal male** | ***Ceasia* focal female** | **Rival male** | **n** |
| --- | --- | --- | --- |
| Allopatric *E. spectabile* | Allopatric *E. spectabile* | Allopatric *E. spectabile* | 12 |
| Allopatric *E. pulchellum* | Allopatric *E. pulchellum* | Allopatric *E. pulchellum* | 12 |

**B. Male competition trials with *E. caeruleum* rival male**
| ***Ceasia* focal male** | ***Ceasia* focal female** | **Rival male** | **n** |
| --- | --- | --- | --- |
| Allopatric *E. spectabile* | Allopatric *E. spectabile* | Allopatric *E. caeruleum* | 12 |
| Allopatric *E. pulchellum* | Allopatric *E. pulchellum* | Allopatric *E. caeruleum* | 12 |
| Sympatric *E. spectabile* | Sympatric *E. spectabile* | Sympatric *E. caeruleum* | 12 |

**C. Male competition trials with heterospecific *Ceasia* rival male**
| ***Ceasia* focal male** | ***Ceasia* focal female** | **Rival male** | **n** |
| --- | --- | --- | --- |
| Allopatric *E. spectabile* | Allopatric *E. spectabile* | Allopatric *E. pulchellum* | 12 |
| Allopatric *E. pulchellum* | Allopatric *E. pulchellum* | Allopatric *E. spectabile* | 12 |

Fish were collected with a kick seine in March 2016 and April 2017 and transported back to the laboratory at the University of Illinois at Urbana-Champaign in aerated coolers. Fish were separated into stock aquaria according to population and sex, and were fed daily *ad libitum* with frozen bloodworms. Stock aquaria were maintained at 19° C and fluorescent lighting was provided to mimic the natural photoperiod.

### Comparisons between *Ceasia* and *E. caeruleum*

#### Dichotomous male mate choice assays

In 2016, we first used dichotomous male choice assays to test for a pattern of RCD between *Ceasia* and *E. caeruleum*. In these trials, we placed a single focal male in a test aquarium with a conspecific female and a heterospecific female - *E. caeruleum* (Fig. 2a). This allowed us to determine whether males differ in how they respond to conspecific females versus *E. caeruleum* females. We measured male mate choice for females in three *Ceasia* populations: sympatric *E. spectabile*, allopatric *E. spectabile*, and allopatric *E. pulchellum* (Table 1A). Our prediction was that male mate choice for conspecifics should be high for *E. spectabile* from the sympatric population, and that male mate choice for conspecifics should be low for *E. spectabile* from the allopatric population and for allopatric *E. pulchellum*.

**Figure 2.**
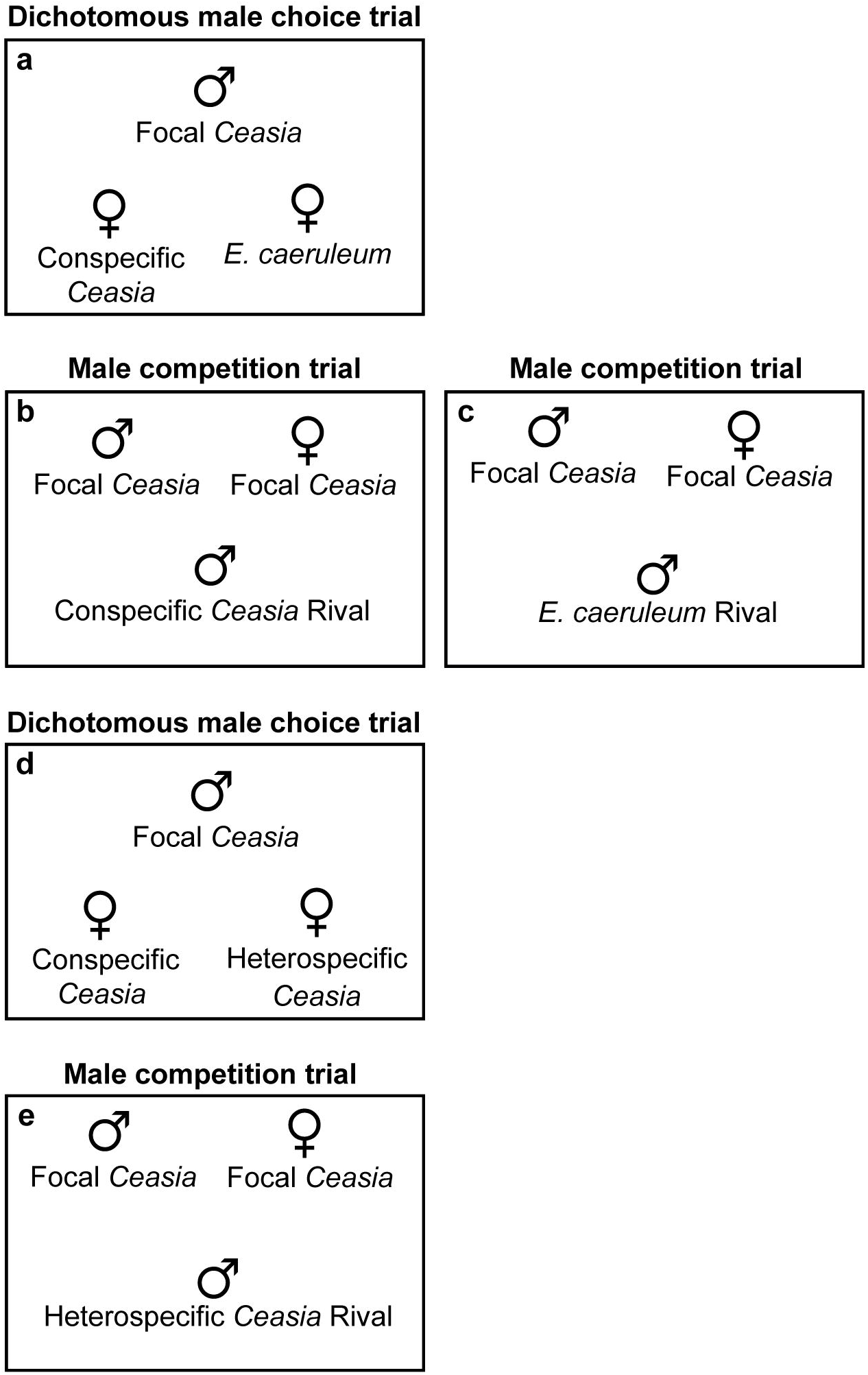
(a-c) Experimental set up for dichotomous male choice trials and male competition trials conducted between *Ceasia* and *E. caeruleum* in 2016. Sympatric *E. spectabile*, allopatric *E. spectabile*, and allopatric *E. pulchellum* acted as focal *Ceasia* in these trials. (d-e) Experimental set up for dichotomous male choice trials and male competition trials conducted between *Ceasia* species in 2017. Allopatric *E. spectabile* and allopatric *E. pulchellum* acted as focal *Ceasia* and as heterospecific *Ceasia* in turn in these trials. In 2017, we did not repeat male competition trials in which a conspecific *Ceasia* acted as the rival male (shown in b). We compared the behavior of individuals in trials with a conspecific *Ceasia* rival male (b) to individuals in trials with an *E. caeruleum* rival male (c). We also compared the behavior of individuals in trials with a conspecific *Ceasia* rival male (b) to individuals in trials with a heterospecific *Ceasia* rival male (e).

Behavioral assays occurred in 38 L test aquaria that were filled with 5 cm of naturally colored aquarium gravel. To minimize disturbance to the fish, test aquaria were covered with black opaque plastic on three sides. Each trial began by placing the three fish being tested into a test aquarium and allowing them to acclimatize for 5 min. The trial then began and lasted 30 min. To avoid exposing fish to chemical cues from fish used in other trials, fish were placed into an aquarium with freshly changed water for each trial. Each 30 min trial was broken up into 60 30-s blocks (Zhou et al. 2015; Moran et al. in press).

We examined male mate choice by measuring focal male pursuit of each female. Previous studies have shown that male pursuit of a female is highly predictive of spawning in *Ceasia* and *E. caeruleum* (Zhou et al. 2015; Moran et al. in press). A male was scored as having pursued a female in a given 30-s block of the trial if he spent a minimum consecutive time of 5-s within one body length of the female. We calculated focal male mate choice as described in Table 3.

**Table 3.**
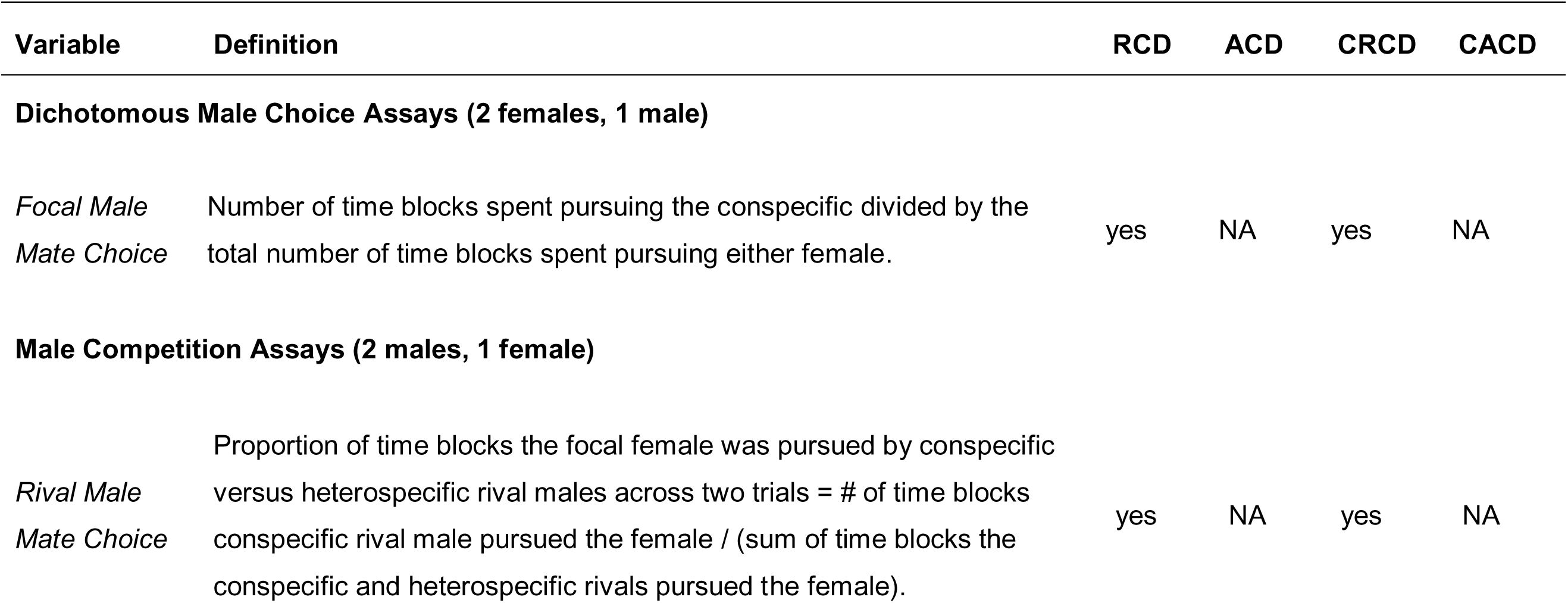
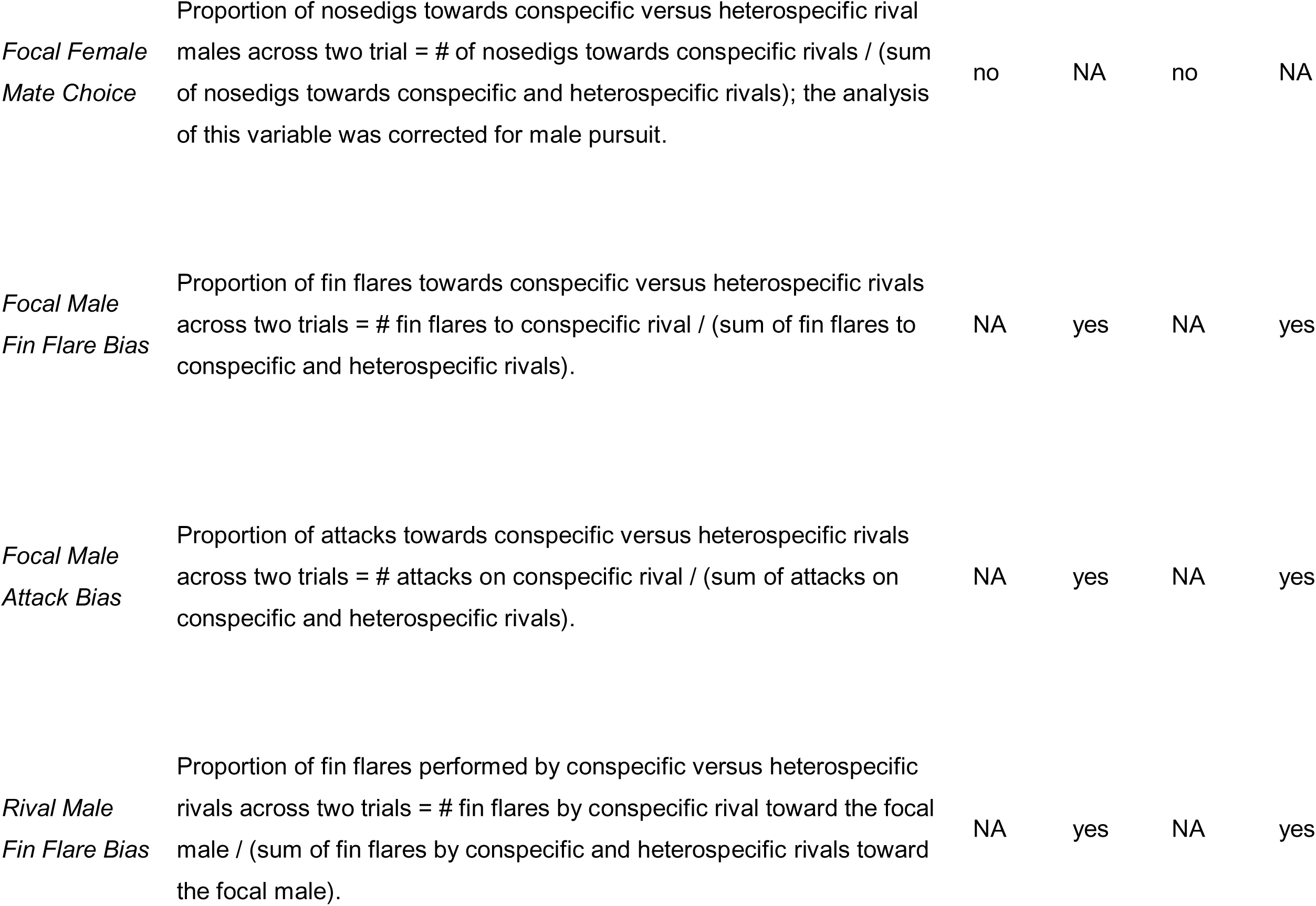
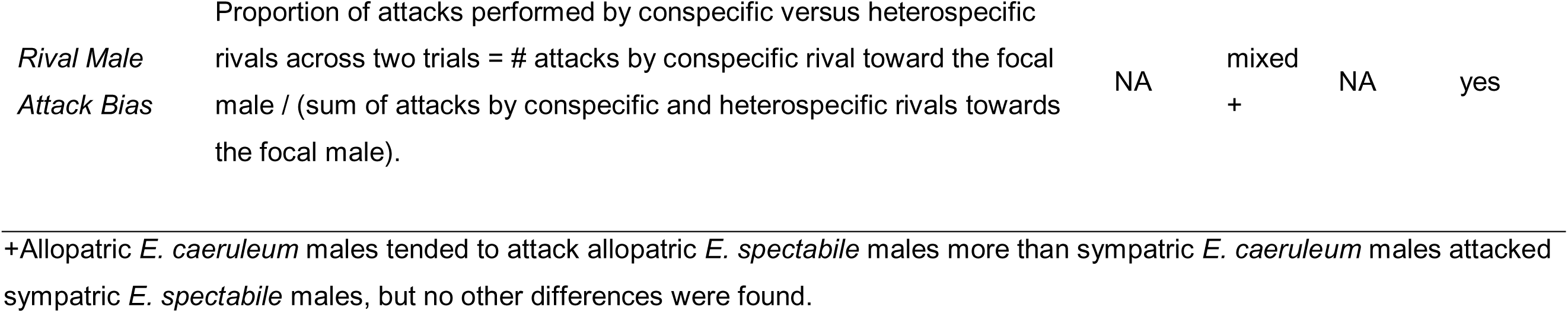
Definition of the behavioral variables measured in dichotomous male mate choice assays and male competition assays. We also indicate whether each behavioral variable showed a pattern that was consistent with our predictions for RCD, ACD, CRCD, and CACD (i.e., lack of discrimination against heterospecifics in allopatric populations, and high discrimination against heterospecifics in sympatric populations). NA denotes that the behavioral variable was not applicable to testing the predictions for that type of character displacement.

For both the dichotomous male choice assays and the male competition assays (see below), analyses were performed using proportional data (i.e., the behavioral variables described in Table 3) that varied from 0 to 1. A score of 1 would indicate only conspecific interactions occurred, 0.5 would indicate an equal number of interactions between conspecifics and heterospecifics, and 0 would indicate only heterospecific interactions occurred. However, for ease of interpretation, we graphed the raw number of behaviors observed.

We used analysis of variance (ANOVA) to determine whether focal male mate choice differed among the three focal *Ceasia* populations (i.e., sympatric *E. spectabile*, allopatric *E. spectabile*, and allopatric *E. pulchellum*) in the dichotomous male choice trials. We included focal male mate choice as the dependent variable, and focal *Ceasia* population as the independent variable. We then used post-hoc t-tests to directly compare populations. We also asked whether focal male mate choice differed from a null expectation of 0.5 (equal amounts of time spent with each female) in each focal *Ceasia* population using sample t-tests.

#### Male competition assays

We conducted a second type of assay in 2016 in which males could compete with one another. This assay allowed us to look for patterns of RCD and ACD between *Ceasia* and *E. caeruleum*. We conducted the 2016 male competition assay using the same *Ceasia* and *E. caeruleum* study populations as in the 2016 dichotomous male choice assay (Table 2). Male competition trials included a focal male and focal female *Ceasia* pair, and a rival male that was either (a) conspecific to the focal *Ceasia* pair or (b) an *E. caeruleum.* Each focal male and focal female *Ceasia* pair was observed twice: once in a trial where the rival male was a conspecific *Ceasia* (Fig. 2b; Table 2A), and once in a trial where the rival male was an *E. caeruleum* (Fig. 2c; Table 2B). Due to low collection numbers, some allopatric *E. caeruleum* males were used twice (i.e, once as a rival male in a trial with allopatric *E. spectabile* and once as a rival male in a trial with allopatric *E. pulchellum*), but never more than once on the same day or with the same *Ceasia* population.

The male competition assay allowed us ask whether sympatric male *Ceasia* and *E. caeruleum* were more likely to discriminate against heterospecific males in a competitive context compared to allopatric male *Ceasia* and *E. caeruleum*. We also asked whether sympatric males showed higher levels of discrimination against heterospecific females compared to allopatric males when they could simply choose whether or not to pursue a female (i.e., a no-choice situation). Additionally, this assay allowed us to examine whether female mate preference for conspecific males versus *E. caeruleum* males was present, and whether it differed among allopatric and sympatric populations.

To measure the aggressive response of both males (focal and rival) towards the other male in each trial, we recorded the number of aggressive behaviors (i.e., fin flares and attacks) performed by each male. We calculated four behavioral variables to quantify male discrimination against heterospecific males: focal male fin flare bias, focal male attack bias, rival male fin flare bias, and rival male attack bias (see Table 3 for variable calculations).

To measure male mate preference in the male competition trials, trials were split into 60 30-s blocks (as in the dichotomous male choice trials) and we counted the number of 30-s blocks in which each male pursued the female in each trial. We calculated rival male mate choice as described in Table 3. We did not measure focal male mate choice in the male competition trials, as focal males were always paired with a conspecific female in these trials. Unlike the dichotomous male choice assay, the male competition assay examines male mate preference in the presence of a male competitor (which is closer to what a male would most often face in nature during the spawning season). Additionally, the male competition assay considers the preference of male *E. caeruleum* for *Ceasia* females, whereas the dichotomous choice trials only considered the preference of *Ceasia* males for conspecific females versus heterospecific *E. caeruleum* females. As male mate preference has been previously shown to be important in maintaining species boundaries in these species (Zhou et al. 2015; Moran et al. in press), examining mate preference from both the perspective of male *Ceasia* and male *E. caeruleum* is critical to estimating behavioral isolation between species.

Finally, we measured focal female mate choice by counting the number of nosedigs a female performed towards the rival male in each trial (see Table 3). Females typically perform nosedigs directly before spawning, and this behavior is often used to measure female mating preferences in darters (Fuller 2003; Williams and Mendelson 2011; Zhou et al. 2015; Zhou and Fuller 2016).

To test for a pattern of ACD between *Ceasia* and *E. caeruleum*, we asked whether the behavioral variables that we used to measure male aggression differed among sympatric and allopatric populations. To examine focal male behavior, we conducted two separate ANOVAs with focal male fin flare bias and focal male attack bias as the dependent variables, and focal *Ceasia* population as the independent variable in both analyses. Similarly, to examine rival male behavior, we conducted ANOVAs with rival male fin flare bias and rival male attack bias as dependent variables, and focal *Ceasia* population as the independent variable. Additionally, we performed post-hoc analyses using two-sample t-tests to make direct comparisons between focal *Ceasia* populations.

To test for a pattern of RCD in male mate preferences, we asked whether male mate choice differed among sympatric and allopatric focal *Ceasia* populations. We conducted an ANOVA with male mate choice as the dependent variable, and focal *Ceasia* population as the independent variable, followed by post-hoc two-sample t-tests between populations.

To test for a pattern of RCD via increased female discrimination against heterospecific males in sympatry, we asked whether focal female mate choice differed among the three focal *Ceasia* populations examined using ANCOVA. The model included focal female mate choice as the dependent variable and focal *Ceasia* population as the independent variable. We also included the proportion of time that conspecific rival males pursued the *Ceasia* focal female as a covariate in the analysis, since previous studies have shown that male pursuit predicts female nosedigs and spawning (Zhou et al. 2015; Moran et al. in press). As the goal of this analysis was to test for a pattern of increased female preference for conspecific mates in sympatry compared to allopatry (i.e., RCD in female mate preference), and previous studies have indicated that females do not exert preferences among males within or among species (Pyron 1995; Fuller 2003; Zhou et al. 2015; Moran et al. in press), we also used ANCOVA to test for female mate choice between conspecific and *E. caeruleum* rival males within each focal *Ceasia* population. For these within-population analyses, the number of nosedigs the focal females directed towards each rival male was the independent variable, the rival male’s identity (conspecific or *E. caeruleum*) was the dependent variable, and the proportion of time the rival male spent in pursuit of the female was included as a covariate.

### Comparisons between Ceasia species

#### Dichotomous male choice assays

In 2017, we examined behavioral isolation between allopatric *Ceasia* species. We asked whether allopatric *Ceasia* males were able to discriminate between conspecific females and allopatric heterospecific *Ceasia* females. The hypothesis that CRCD is occurring within *Ceasia* predicts that male discrimination against heterospecific females should be low between the two allopatric *Ceasia* species (as neither occur in sympatry with *E. caeruleum*, and thus do not experience reinforcement). To test this, we compared the allopatric *E. spectabile* and allopatric *E. pulchellum* study populations that were used in the 2016 comparisons between *Ceasia* and *E. caeruleum.*

We performed dichotomous male choice assays as described above for the *Ceasia* and *E. caeruleum* comparisons conducted in 2016, but with a heterospecific *Ceasia* female in place of the *E. caeruleum* female (Fig. 2d). We performed trials in which allopatric *E. spectabile* acted as the focal male and conspecific female, with *E. pulchellum* as the heterospecific *Ceasia* female, and vice versa (Table 1B). We asked whether male preference for conspecifics differed among the two allopatric *Ceasia* populations. As this set of trials only compared allopatric populations, we expected to find no significant difference between the two *Ceasia* focal populations in focal male mate choice (Table 3). To compare male mate choice between populations, we included focal male mate choice as the dependent variable, and focal *Ceasia* population (allopatric *E. spectabile* or allopatric *E. pulchellum*) as the independent variable. We also tested whether focal male mate choice for the conspecific female differed from a null expectation of 0.5 (equal amounts of time spent with each female).

#### Male competition assays

We also conducted male competition assays between the two allopatric *Ceasia* species in 2017. A previous study by Moran et al. (in press) found high levels of male-driven behavioral isolation between *Ceasia* species that occur in sympatry with *E. caeruleum*. Our hypothesis that CRCD is present in *Ceasia* predicts low levels of mate discrimination against heterospecific *Ceasia* in species that occur allopatrically from *E. caeruleum*. Additionally, our hypothesis that CACD is present in *Ceasia* predicts low levels of male competitive discrimination against heterospecific *Ceasia* males in species that occur allopatrically from *E. caeruleum*. To test these hypotheses, we performed male competition assays as described above for the trials examining interactions between *Ceasia* and *E. caeruleum* conducted in 2016, but using a heterospecific *Ceasia* rival male in place of the *E. caeruleum* rival male (Fig. 2e). In these trials, we paired allopatric *E. spectabile* with allopatric *E. pulchellum* (Table 2C). We performed trials in which each of these allopatric *Ceasia* species acted as the focal pair and as the heterospecific rival male. We did not repeat trials conducted in 2016 in which each of these *Ceasia* species were paired with a conspecific rival male.

We measured male aggression, male mate choice, and female mate choice as described in Table 3. We conducted ANOVAs as described above for the 2016 male competition trials that paired *Ceasia* with *E. caeruleum*, but with heterospecific *Ceasia* in place of *E. caeruleum.* As the analyses for these 2017 trials compared discrimination against heterospecifics in two allopatric *Ceasia* populations, our prediction for CRCD and CACD was that there would be no significant differences between these two populations (i.e., both allopatric *Ceasia* populations should show low levels of discrimination against one another).

### Behavioral isolation indices

We used the male aggression, male mate choice, and female mate choice data from the male competition assays to calculate three behavioral isolation indices following Moran et al. (in press). Behavioral isolation indices were calculated individually for each trial and then averaged across all replicates within each species comparison. These indices allowed for a comparison among species pairs (i.e., for each *Ceasia - E. caeruleum* and *Ceasia - Ceasia* comparison) of levels of discrimination against heterospecifics. Indices range from -1 (complete preference for heterospecifics) to 1 (complete preference for conspecifics), with 0 indicating no preference for conspecifics versus heterospecifics (Stalker 1942; Martin and Mendelson 2016; Moran et al. in press).

We calculated male aggression (MA) indices for each species pair as:

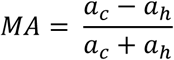

where a_c_ and a_h_ represent the combined number of fin flares and attacks performed between conspecific males and between heterospecific males, respectively.

We calculated male choice (MC) indices as:

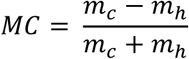

where m_c_ and m_h_ represent the proportion of time in each trial that conspecific males and heterospecific males spent pursuing the *Ceasia* female.

As previous studies have indicated that male pursuit of a female is highly correlated with female nosedigs (a measure of female mating preference), female choice (FC) indices controlled for male pursuit of the female. We calculated the FC indices as:

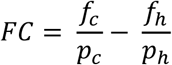

where f_c_ and f_h_ represent the number of nosedigs females performed towards conspecific males and towards heterospecific males, respectively. p_c_ and p_h_ represent the number of 30-s blocks in which conspecific males and heterospecific males were scored as having pursued the female during a trial, respectively.

## Results

### Reproductive Character Displacement between *Ceasia* and *E. caeruleum*

There was a clear pattern of RCD between *Ceasia* and *E. caeruleum* in the dichotomous male choice trials, due to increased male mate discrimination against *E. caeruleum* females in sympatry. Focal male mate choice differed among sympatric and allopatric *Ceasia* populations, but not between allopatric *Ceasia* populations (Table 4). Sympatric *E. spectabile* focal males showed 2X higher levels of discrimination against sympatric *E. caeruleum* females than did allopatric *E. spectabile* males (Fig. 3a). Sympatric *E. spectabile* male mate choice was much greater than the null expectation of 0.5 (mean ± SE: 0.97 ± 0.01; one-sample t-test: t_11_=51.58, p<0.00001). Conversely, male mate choice did not differ from 0.5 in the allopatric *Ceasia* males (Fig. 3b,c; allopatric *E. spectabile* mean ± SE: 0.51 ± 0.04; one-sample t-test: t_11_=0.17, p=0.87; *E. pulchellum* mean ± SE: 0.53 ± 0.05; one-sample t-test: t_11_=0.60, p=0.56).

**Table 4.**
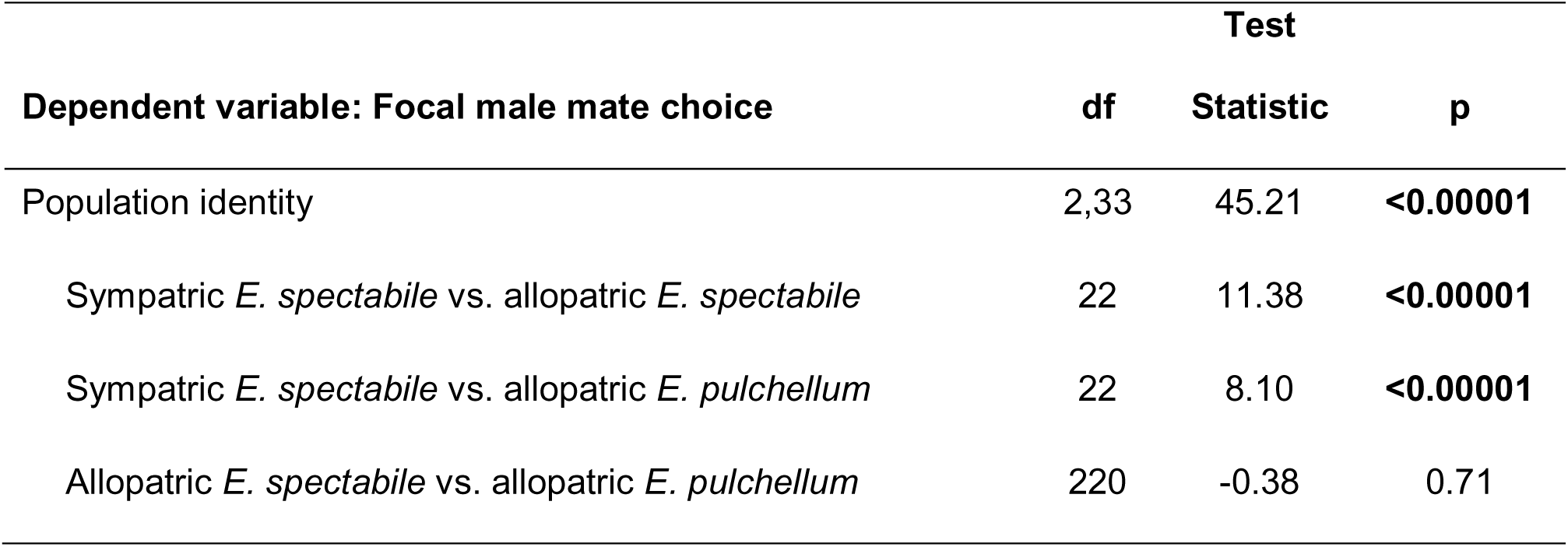
Results of ANOVA on focal male mate choice between conspecific females and *E. caeruleum* females in dichotomous male choice male trials. We asked focal male mate choice differed among focal *Ceasia* populations (sympatric *E. spectabile*, allopatric *E. spectabile*, and allopatric *E. pulchellum*). Pairwise post-hoc t-test results are also shown for the analysis.

**Figure 3.**
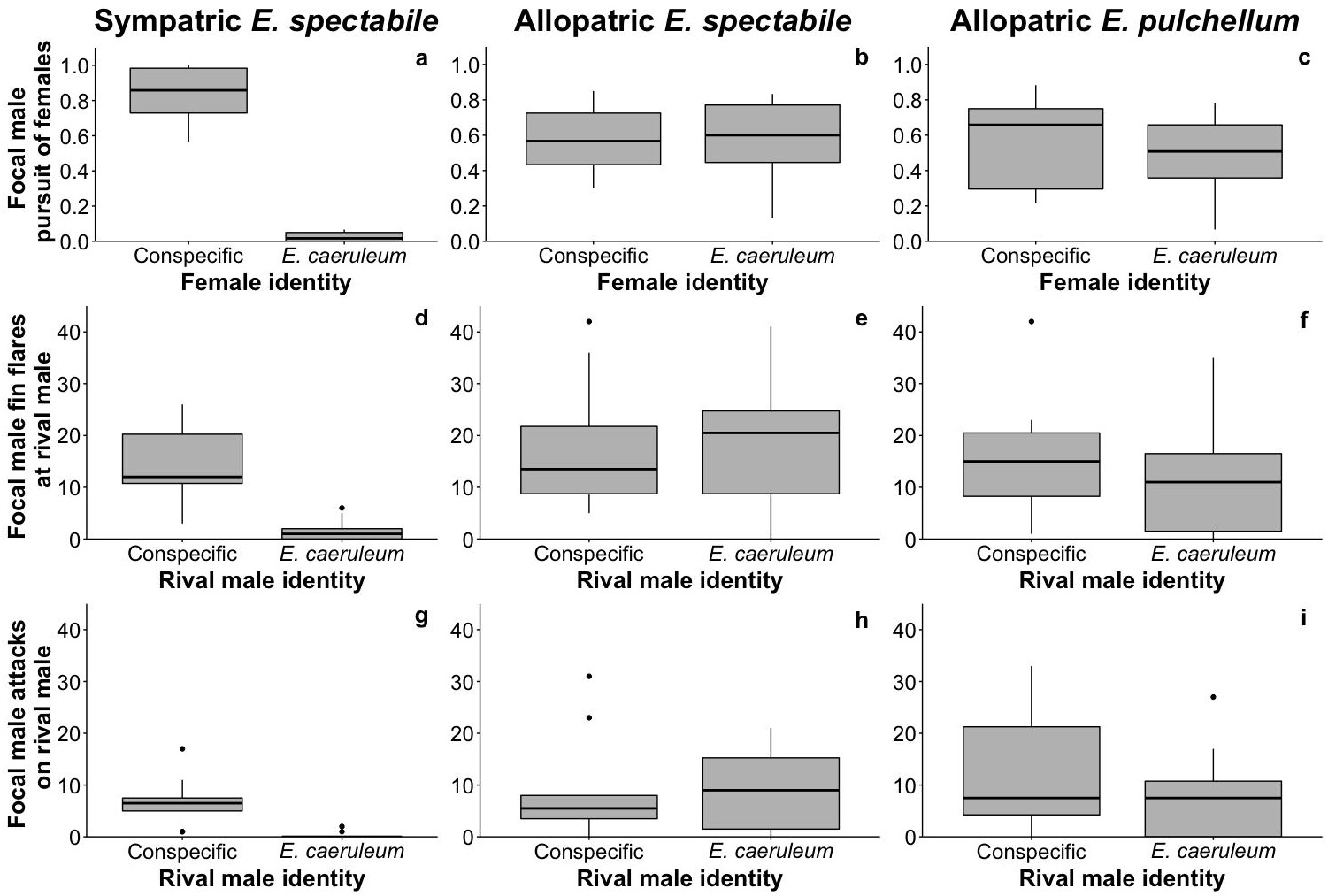
Focal male behavior in trials that paired *Ceasia* with *E. caeruleum.* Columns from left to right show results for trials with sympatric *E. spectabile*, allopatric *E. spectabile*, and allopatric *E. pulchellum* as the focal *Ceasia*, respectively. (a-c) Proportion of time focal males spent in pursuit of conspecific versus *E. caeruleum* female in dichotomous choice trials. (d-f) Number of focal male fin flares directed at conspecific versus *E. caeruleum* rival males in male competition trials. (g-i) Number of focal male attacks directed at conspecific versus *E. caeruleum* rival males in male competition trials.

RCD in male mate choice (i.e., increased discrimination against heterospecific females in sympatric populations) was also indicated in the male competition trials that considered male *E. caeruleum* mate choice. Rival male mate choice differed significantly among sympatric and allopatric *Ceasia* populations, but not between allopatric *Ceasia* populations (Table S2). In sympatric trials, conspecific rival males were much more likely to pursue the focal female *Ceasia* compared to *E. caeruleum* rival males (Fig. S1a). In allopatric trials, conspecific and *E. caeruleum* rival males and spent roughly the same amount of time pursuing the focal female *Ceasia*. Hence, sympatric *E. caeruleum* males discriminated against sympatric *E. spectabile* females, but allopatric *E. caeruleum* males did not discriminate against allopatric *E. spectabile* or allopatric *E. pulchellum* females.

Unlike male *Ceasia*, female *Ceasia* mating preference for conspecific versus *E. caeruleum* males did not differ among the focal *Ceasia* populations. When male pursuit was included as a covariate in the analysis, female mate choice did not differ among *Ceasia* populations (Table 5). This was due to females not exerting any preference for conspecific males over *E. caeruleum* males across all three populations, regardless of sympatry with *E. caeruleum* (Table S3).

**Table 5.** Results ANCOVA examining focal female mate choice between conspecific rival males and *E. caeruleum* rival males in male competition trials. We asked whether female mate choice differed among focal *Ceasia* populations (sympatric *E. spectabile*, allopatric *E. spectabile*, and allopatric *E. pulchellum*). Male pursuit of the female was included as a covariate in the analysis.

| Dependent variable: Focal female mate choice | Test |  |  |
| --- | --- | --- | --- |
|  | df | Statistic | p |
| Population identity | 2,32 | 0.09 | 0.92 |
| Male pursuit | 1,32 | 0.74 | 0.40 |

### Agonistic Character Displacement between *Ceasia* and *E. caeruleum*

The male competition trials revealed a pattern consistent with ACD between *Ceasia* and *E. caeruleum*, due to increased male discrimination against heterospecific competitors in sympatry. Focal male fin flare bias and focal male attack bias differed significantly among sympatric versus allopatric *Ceasia* populations, but did not differ between allopatric *Ceasia* populations (Table 6). Sympatric *E. spectabile* focal males directed 9x more fin flares towards conspecific rival males compared to *E. caeruleum* rival males (Fig. 3d). Similarly, sympatric *E. spectabile* focal males attacked conspecific rival males 6x more than they attacked sympatric *E. caeruleum* rival males (Fig. 3g). On average, both allopatric *E. spectabile* and allopatric *E. pulchellum* focal males directed an equal number of fin flares (Fig. 3e,f) and attacks (Fig. 3h,i) towards conspecific rival males and allopatric *E. caeruleum* rival males.

**Table 6.** Results of ANOVA on focal male *Ceasia* fin flares and attacks at rival males in male competition trials. We asked whether focal male fin flare bias and focal male attack bias differed among focal *Ceasia* populations (sympatric *E. spectabile*, allopatric *E. spectabile*, and allopatric *E. pulchellum*). Pairwise post-hoc t-test results are also shown for both analyses.

| Dependent variable: Focal male fin flare bias | Test |  |  |
| --- | --- | --- | --- |
|  | df | Statistic | p |
| Population identity | 2,33 | 8.34 | **0.0012** |
| Sympatric *E. spectabile* vs. allopatric *E. spectabile* | 22 | 5.28 | **<0.0001** |
| Sympatric *E. spectabile* vs. allopatric *E. pulchellum* | 22 | 2.85 | **0.0093** |
| Allopatric *E. spectabile* vs. allopatric *E. pulchellum* | 22 | -0.84 | 0.41 |

| Dependent variable: Focal male attack bias | Test |  |  |
| --- | --- | --- | --- |
|  | df | Statistic | p |
| Population identity | 2,33 | 9.12 | **<0.001** |
| Sympatric *E. spectabile* vs. allopatric *E. spectabile* | 22 | 4.53 | **0.0002** |
| Sympatric *E. spectabile* vs. allopatric *E. pulchellum* | 22 | 3.82 | **<0.001** |
| Allopatric *E. spectabile* vs. allopatric *E. pulchellum* | 22 | -0.65 | 0.52 |

Rival male fin flare bias also showed a pattern consistent with ACD and was similar to what was found with focal males (Table S4). Sympatric *E. caeruleum* rival males were much less likely to flare their fins towards *E. spectabile* males than were allopatric *E. caeruleum* rival males (Fig. S1d-f). Conversely, rival male attack bias did not differ among trials with sympatric versus allopatric focal *Ceasia* (Table S4). Both sympatric and allopatric *E. caeruleum* directed a low number of attacks towards focal male *Ceasia* (Fig. S1g-i). Thus, while focal males in allopatric *Ceasia* populations failed to discriminate against allopatric *E. caeruleum* males, allopatric *E. caeruleum* rival males did discriminate somewhat against allopatric *Ceasia* males.

### Mate discrimination between allopatric *E. spectabile* and allopatric *E. pulchellum*

As expected, both allopatric populations of *Ceasia* failed to discriminate against heterospecific *Ceasia* mates. In dichotomous male choice trials, focal male mate choice did not differ between allopatric *E. spectabile* and allopatric *E. pulchellum* (F_1,22_ = 0.29; p = 0.60; Fig. S2a,b). Allopatric *E. spectabile* focal male mate choice did not differ from a null expectation of 0.5 (mean ± SE: 0.42±0.04; one-sample t-test: t_11_ = -1.94, p = 0.08;). The same pattern was observed for allopatric *E. pulchellum* focal male mate choice (mean ± SE: 0.45±0.04; one-sample t-test: t_11_ = -1.28, p = 0.23). Similarly, rival male mate choice did not differ between allopatric *E. spectabile* and allopatric *E. pulchellum* in the male competition trials (F_1,22_ = 0.12; p = 0.73; Fig. S3).

Additionally, focal female mate choice did not differ between allopatric *E. spectabile* and allopatric *E. pulchellum*, and these preferences did not differ from 0.5 (Table S5). There was no significant difference in the proportion of female nosedigs towards rival males as function of their identity (conspecific or heterospecific *Ceasia*) when we controlled for the proportion of time each male pursued the female (Table S6).

### Competitor discrimination between allopatric *E. spectabile* and allopatric *E. pulchellum*

Consistent with our prediction, neither of the populations of allopatric *Ceasia* discriminated against heterospecific *Ceasia* male competitors. Focal male fin flare bias did not differ between allopatric *E. spectabile* and allopatric *E. pulchellum* (F1,22 = 1.79; p = 0.19; Fig. S2c,d), nor did focal male attack bias (F1,22 = 0.84; p = 0.37; Fig. S2e,f).

Allopatric *Ceasia* males also failed to discriminate against heterospecific *Ceasia* males when they acted as the rival male. In the trials where allopatric *E. pulchellum* were the focal males, conspecific rival males and allopatric *E. spectabile* (heterospecific *Ceasia*) rival males directed a similar number of fin flares towards focal males (Fig. S3d). However, in trials where allopatric *E. spectabile* were the focal males, allopatric *E. pulchellum* (heterospecific *Ceasia*) rival males directed more fin flares towards the focal males compared to the conspecific rival males (Fig. S3c). This resulted in a significant difference in rival male fin flare bias between the two allopatric *Ceasia* populations (F1,22 = 5.79; p = 0.025; Fig. S3), despite the pattern being consistent with our prediction for CACD. Rival male attack bias did not differ between trials with allopatric *E. spectabile* versus allopatric *E. pulchellum* as the focal *Ceasia* (F1,22 = 0.10; p = 0.75; Fig. S3).

### Is there a pattern consistent with RCD and CRCD across *Ceasia*?

To examine patterns of character displacement across *Ceasia*, we compared the behavioral isolation indices we calculated in this study with behavioral isolation indices calculated by Moran et al. (in press) (Tables 7, S7; Fig. 4). Behavioral isolation was measured between the same populations of sympatric *E. spectabile* and sympatric *E. caeruleum* in the present study and in Moran et al (in press; Table S7). Calculations of behavioral isolation indices for the pairing of sympatric *E. spectabile* with sympatric *E. caeruleum* did not differ between this study and Moran et al (in press) (two-sample t-tests; MA: t_38_= 0.31, p=0.76; MC: t_18_= -1.44, p=0.17; FC: t_18_= -0.98, p=0.34). Thus, the MA, MC, and FC indices presented here for the pairing of sympatric *E. spectabile* with sympatric *E. caeruleum* (Table 7; Fig. 4) were calculated by pooling the behavioral isolation data from this study with behavioral isolation data from Moran et al. (in press).

**Table 7.** Behavioral isolation indices (mean ± standard error) for male aggression (MA), male choice (MC), and female choice (FC). For each species pair, the *Ceasia* species that acted as the focal *Ceasia* in behavioral trials is listed first, followed by the species that it was observed with (a heterospecific *Ceasia* or *E. caeruleum*).

| Geography | Pairing | Species | MA | MC | FC |
| --- | --- | --- | --- | --- | --- |
| Allopatric | *Ceasia* - *Ceasia* | *E. spectabile* - *E. pulchellum* | -0.01±0.07 | 0.11±0.07 | 0.01±0.02 |
| Sympatric | *Ceasia* - *Ceasia* | *E. fragi* - *E. uniporum* * | 0.38±0.08 | 0.31±0.07 | 0.01±0.01 |
| Sympatric | *Ceasia* - *Ceasia* | *E. fragi* - *E. burri* * | 0.50±0.06 | 0.30±0.07 | 0.02±0.01 |
| Sympatric | *Ceasia* - *Ceasia* | *E. fragi* - *E. spectabile* * | 0.35±0.06 | 0.34±0.10 | 0.01±0.02 |
| Allopatric | *Ceasia* - *E. caeruleum* | *E. spectabile* - *E. caeruleum* | 0.09±0.09 | 0.22±0.12 | -0.16±0.16 |
| Allopatric | *Ceasia* - *E. caeruleum* | *E. pulchellum* - *E. caeruleum* | 0.30±0.12 | 0.25±0.12 | 0.01±0.02 |
| Sympatric | *Ceasia* - *E. caeruleum* | *E. fragi* - *E. caeruleum* * | 0.80±0.05 | 0.76±0.06 | 0.01±0.04 |
| Sympatric | *Ceasia* - *E. caeruleum* | *E. uniporum* - *E. caeruleum* * | 0.82±0.06 | 0.70±0.09 | -0.11±0.13 |
| Sympatric | *Ceasia* - *E. caeruleum* | *E. burri* - *E. caeruleum* * | 0.92±0.03 | 0.66±0.08 | -0.05±0.05 |
| Sympatric | *Ceasia* - *E. caeruleum* | *E. spectabile* - *E. caeruleum* ** | 0.85±0.05 | 0.84±0.06 | 0.03±0.02 |

\*From Moran et al. (in press).
\*\*Calculated using data from the present study combined with data from Moran et al. (in press).

**Figure 4.**
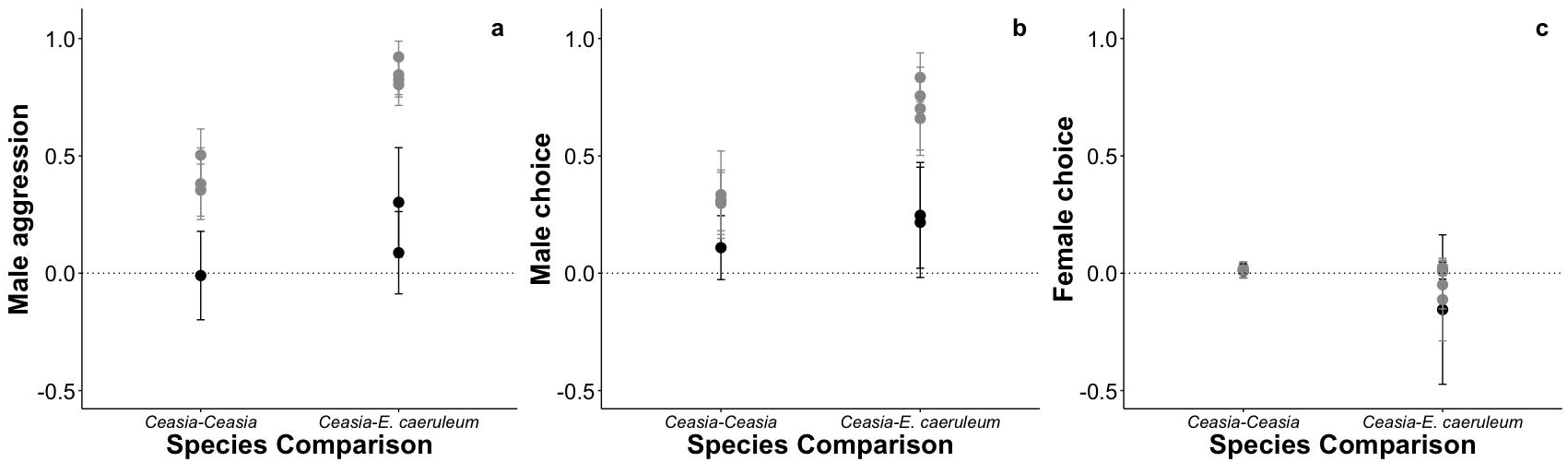
Behavioral isolation indices (with 95% confidence intervals) for (a) male aggression, (b) male choice, and (c) female choice. Each point represents a comparison between two *Ceasia* species (*Ceasia - Ceasia*) or between a *Ceasia* species and *E. caeruleum* (*Ceasia - E. caeruleum*). Allopatric comparisons (i.e., comparisons including *Ceasia* that do not co-occur with *E. caeruleum*) are shown in black. Sympatric comparisons (i.e., comparisons including *Ceasia* that do co-occur with *E. caeruleum*) are shown in gray.

RCD predicts higher levels of discrimination against heterospecific mates (i.e., higher MC and/or FC indices) in sympatric compared to allopatric *Ceasia - E. caeruleum* pairs. CRCD predicts increased levels of discrimination against heterospecific mates (i.e., higher MC and/or FC indices) in sympatric compared to allopatric *Ceasia* - *Ceasia* pairs. In other words, *Ceasia* that are sympatric with *E. caeruleum* (but allopatric to one another) should have increased behavioral isolation.

We observed a pattern consistent with RCD and CRCD. MC behavioral isolation indices were consistently higher between sympatric species pairs compared to allopatric species pairs, both within the *Ceasia* - *E. caeruleum* comparisons and within the *Ceasia* - *Ceasia* comparisons (Table 7; Fig. 4b). The difference between sympatry and allopatry was more pronounced in *Ceasia* - *E. caeruleum* comparisons.

There was no pattern among FC indices as a function of sympatry with *E. caeruleum* (Table 7; Fig. 4c). This was due to females not exerting any detectable mating preferences for conspecific males.

### Is there a pattern consistent with ACD and CACD across *Ceasia*?

ACD predicts higher levels of male discrimination against heterospecific rival males (i.e., higher MA indices) in sympatric compared to allopatric *Ceasia - E. caeruleum* pairs. CACD predicts increased levels of male discrimination against heterospecific rival males (i.e., higher MA indices) in sympatric compared to allopatric *Ceasia* - *Ceasia* pairs.

We observed a pattern consistent with ACD and CACD. MA behavioral isolation indices were consistently higher between sympatric species pairs compared to allopatric species pairs, both within the *Ceasia* - *E. caeruleum* comparisons and within the *Ceasia* - *Ceasia* comparisons (Table 7; Fig. 4a). As with MC, the difference between sympatry and allopatry was more pronounced in *Ceasia* - *E. caeruleum* comparisons.

## Discussion

The results of this study suggest that (1) reinforcement has occurred multiple times between *Ceasia* and *E. caeruleum* throughout their range of sympatry, and (2) cascading effects of reinforcement between *Ceasia* and *E. caeruleum* has incidentally contributed to allopatric speciation within the *Ceasia* clade. Although theory predicts that cascading effects of reinforcement can lead to allopatric speciation (McPeek and Gavrilets 2006; Pfennig and Ryan 2006), the majority of empirical studies that have examined cascade reinforcement to date have tested behavioral preferences among closely related populations within species. In addition, many other studies have compared populations within species where there is behavioral isolation between populations that are allopatric versus sympatric with another species (Nosil et al. 2003; Lemmon 2009; Hopkins et al. 2014; Kozak et al. 2015; Comeault and Matute 2016; Comeault et al. 2016). The implication with these studies is that reinforcement changes preferences and target traits in such a way that increases behavioral isolation between sympatric and allopatric populations (i.e., “sympatry-allopatry effects”). Here, there are high levels of behavioral isolation between *Ceasia* species that have independently undergone reinforcement with *E. caeruleum.* This suggests that different species-specific preferences and traits have evolved in *Ceasia* species that are sympatric with *E. caeruleum* (i.e., “convergent-sympatry effects”). This study provides an important example of how cascade reinforcement can cause high levels of behavioral isolation to evolve between closely related populations and result in allopatric speciation.

We observed a striking pattern of RCD and ACD between *Ceasia* and *E. caeruleum*, primarily driven by male *Ceasia* behavior. Sympatric *E. spectabile* males strongly discriminated against *E. caeruleum* female mates and male competitors, but allopatric *E. spectabile* and allopatric *E. pulchellum* males did not discriminate against allopatric *E. caeruleum* of either sex. As expected, allopatric *E. spectabile* and allopatric *E. pulchellum* also did not discriminate against one another in a mating or fighting context. Furthermore, our results agree with several previous studies in this system that have failed to detect female mate preference for conspecific versus heterospecific males in sympatric or allopatric populations of *Ceasia*. Thus, it appears that males play an important role in maintaining species boundaries in these species.

There is also a pattern consistent with our predictions for RCD and ACD in male *E. caeruleum*, but the pattern of ACD was less striking in *E. caeruleum* males compared to *Ceasia* males. Sympatric *E. caeruleum* discriminated against sympatric *E. spectabile* females and males. However, *E. caeruleum* males did not show as low a level of discrimination against *Ceasia* males in allopatry. We hypothesize that the discrimination against *Ceasia* males demonstrated by allopatric *E. caeruleum* males may be due to differences in the level of gene flow experienced by *Ceasia* versus *E. caeruleum*. Patterns of character displacement due to reinforcement are more likely to be maintained over time (and to lead to cascading effects among populations within species) when gene flow is low among populations (Hoskin et al. 2005; Lemmon 2009; Kozak et al. 2015; Yukilevich and Aoki 2016). *Ceasia* and *E. caeruleum* both occur in small headwater streams, but *E. caeruleum* can also inhabit larger order streams and rivers (Page 1983; Page and Burr 2011), leading to more opportunities for gene flow among populations (Echelle et al. 1975, 1976). Indeed, population genetic analyses of four species of *Ceasia* and *E. caeruleum* found increased heterozygosity and higher levels nucleotide diversity are present within *E. caeruleum* relative to the *Ceasia* species, indicating lower levels of gene flow in species of *Ceasia*. This key difference in the biology of *Ceasia* and *E. caeruleum* may explain why *E. caeruleum* has not diversified to the extent that *Ceasia* has, despite being similarly widespread.

Our results together with the results of a recent study by Moran et al. (in press) support the hypothesis that reinforcement has occurred multiple times between *Ceasia* and *E. caeruleum.* Sympatric *Ceasia* species have consistently shown almost complete levels of behavioral isolation with their respective sympatric populations of *E. caeruleum*, but allopatric *Ceasia* do not show any such discrimination (this study; Moran et al. in press; Zhou and Fuller 2014). The observed pattern of RCD and ACD across *Ceasia* together evidence of high levels of postzygotic isolation between *Ceasia* and *E. caeruleum* (Zhou 2014; R. Moran unpubl. data) suggests that reinforcement is responsible for driving behavioral isolation between *Ceasia* and *E. caeruleum* in sympatry.

We also observed a pattern consistent with CRCD and CACD across *Ceasia*. Species of *Ceasia* that occur in sympatry with *E. caeruleum* show surprisingly high levels of male discrimination against heterospecific *Ceasia* mates and competitors, but no such discrimination is present among *Ceasia* that occur in allopatry from *E. caeruleum*. This observation together with the evidence for reinforcement between *Ceasia* and *E. caeruleum* suggest that cascade reinforcement is occurring within *Ceasia*. Cascade reinforcement may occur if slightly different mating traits (signals and/or preferences) arise in different populations across the range of a species experiencing reinforcement with a heterospecific. Theory predicts that cascade reinforcement can readily occur when gene flow between populations within a species is low (as is the case with organisms that occur in isolated headwater streams, such as darters), and when populations respond to reinforcing selection on mating traits and their underlying loci in unique ways due to stochastic processes (i.e., mutation-order selection; Abbott et al. 2013; Mendelson et al. 2014; Comeault and Matute 2016; Yukilevich and Aoki 2016). Under mutation-order selection, species divergence may occur despite the presence of similar types of ecological and sexual selection. In this way, stochastic variation in response to the same selective pressures (i.e., selection against maladaptive heterospecific interactions in sympatry) can potentially lead to allopatric divergence among populations within species.

This study corroborates the results of several recent studies that have shown that male mate choice and male competition play an important role in driving sympatric and allopatric speciation in darters (Ciccotto et al. 2013; Zhou et al. 2015; Zhou and Fuller 2016; Martin and Mendelson 2016; Moran et al. in press). Multiple studies in several different species of darters have also found little or no female mate preference for conspecific over heterospecific males (Martin and Mendelson 2013, 2016; Ciccotto et al. 2014; Zhou et al. 2015; but see Williams and Mendelson 2010, 2011). Furthermore, although the presence of elaborate male nuptial coloration is most commonly attributed to intersexual selection via female mate preferences (Panhuis et al. 2001), male coloration in darters appears to be under intrasexual selection due to intense male-male competition (Zhou et al. 2015; Martin and Mendelson 2016; Zhou and Fuller 2016). Reproductive and agonistic character displacement can lead to shifts in behavioral response to heterospecifics and the signals used to recognize conspecifics versus heterospecifics in sympatry (Brown and Wilson 1956; Grether et al. 2009). Thus, future studies that examine whether character displacement in male color pattern corresponds to the observed ACD and CACD in male aggressive response to heterospecifics would be of interest.

In conclusion, this study provides empirical evidence of male-driven reinforcement and cascade reinforcement in darters. As far as we are aware, this is the first documented case demonstrating that ACD between species can incidentally lead to CACD among populations within species (or in this case, among closely related species within a clade). Although the clear majority of reinforcement studies to date have focused on the evolution of female mating preferences for males, the results of this study demonstrate that male behavior alone can drive speciation between and within species via reinforcement and cascade reinforcement. This underscores the necessity of considering the behavior of both sexes when evaluating whether reinforcement may be at work in a given system. Finally, this study provides important groundwork for future studies examining the effects of reinforcement on sympatric and allopatric speciation in darters. Further research is needed to explore the extent to which reinforcement has been involved in generating the extraordinary diversity of darters.

## Acknowledgements

We thank M. St. John and L. Mitchem for providing helpful comments on this manuscript. This work was supported by the Cooperative State Research, Education, and Extension Service, US Department of Agriculture, under project number ILLU 875-952, the National Science Foundation (DEB 0953716 and IOS 1701676), and the University of Illinois. The treatment of animals was approved by the Institutional Animal Care and Use Committee under protocol #14097.

